# CrossAffinity: A Sequence-Based Protein-Protein Binding Affinity Prediction Tool Using Cross-Attention Mechanism

**DOI:** 10.64898/2026.02.22.707318

**Authors:** Jia Sheng Guan, Zechen Wang, Yuguang Mu

## Abstract

Protein-protein binding affinity is important for understanding protein interactions within a protein complex and for identifying strong drug-peptide binders to a target protein. Many structure-based models were built previously with reasonable performance. However, such models require protein complex structure as input, which is usually unavailable due to high cost and experimental constraints. To tackle such an issue, the sequence-based CrossAffinity model was constructed in this study, using the cross-attention module to extract contextual information of interacting protein components while separating the protein complex into two distinct parts to predict the protein-protein binding affinity. CrossAffinity managed to outperform all structure-based models and sequence-based models in an S34 test set containing newer protein complex structures and binding affinity values in a timeline while being trained on an older dataset, showing generalisability to new data points. In other test sets, namely S90, S90 subset and S79*, CrossAffinity also managed to outperform all other sequence-based models while maintaining comparable performance to many recently published structure-based models. The acceptable performance and quick inference of CrossAffinity enable it to be deployed in situations requiring the prediction of the binding affinity of many protein complexes that lack structural information.

## Introduction

The discovery of potential novel drugs through purely wet laboratory techniques, such as high-throughput screening (HTS), is a costly venture. For HTS of a million compounds, the cost may range from $500,000 to $1,000,000 [1]. Not to mention, such trial-and-error campaigns do not guarantee any guaranteed outcome [2]. To alleviate the situation, Computer-Aided Drug Design (CADD) employs computer algorithms on chemical and biological data, enabling the simulation and prediction of interactions between a ligand and its drug target, which are typically protein or nucleic acid sequences [2]. As such, CADD serve as a valuable tool to reduce research cost, risk and labour, picking only the most promising drug candidates to be tested in the wet lab. With technological breakthroughs in artificial intelligence (AI) in multiple fields, ranging from image recognition and natural language processing to computer vision, such methodologies can also aid in addressing key challenges in drug development. Through leveraging a vast amount of biological data, AI’s role in drug discovery is multi-faceted, being used to recognise disease biomarkers, identify novel drug targets and predict a drug’s safety and efficacy [3-6].

When it comes to drug design, there are multiple modalities to choose from, each with its specific advantages and disadvantages. Among the variety of drug classes, peptide drugs are a major class. These peptides are made up of amino acid residues, with molecular weights ranging from 0.5 to 5 kDa [7]. Frequently, these drugs function as hormones, growth factors, neurotransmitters, ion channel ligands, or anti-infective agents [8]. They are capable of strong and specific binding to their drug targets, such as cell surface receptors [8]. Compared to commonly used small molecule drugs, therapeutic peptides are more able to inhibit protein-protein interactions (PPIs), which usually have a contact area of 1500 to 3000Å^2^, owing to their greater size [9]. Additionally, peptide drugs have enhanced safety and reduced toxicity and immunogenicity [10]. As such, many research efforts were poured into peptide drug discovery.

For computer-aided peptide drug design, many AI-based models were developed. ProteinMPNN and HighMPNN are two models, capable of designing the sequence of linear and cyclic peptides, respectively [11, 12]. There are also generative diffusion models such as AlphaFold3 and RoseTTAFold, which can predict the binding pose of the peptides and drug target [13, 14]. BindCraft is another model which can both design the peptide sequence and also its binding pose simultaneously when given an input target structure [15]. Aside from generating peptide drug sequences and binding poses, another important use of AI-based models is predicting the binding affinity of the peptide to the drug. To build such a model, there are mainly structure-based or sequence-based models. In structure-based models, the model is given the protein-peptide complex structural information. Examples of such models are ProAffinity-GNN, PPI-Affinity, PRODIGY, PPAP, PPI-Graphomer and SSIF-Affinity [16-21]. ProAffinity-GNN creates inter-molecular and intra-molecular graphs using the 3D protein complex structure and leverage on AttentiveFP architecture to learn and predict protein-protein binding affinity [17, 22]. PPI-Affinity employed 23,040 ProtDCal-generated 3D-structure molecular descriptors as features into a support vector machine [19, 23]. PRODIGY combines interfacial contact and non-interacting surface properties to predict the protein-protein binding affinity [18]. PPAP leveraged information from interfacial contact-based attention and used ESM2 with 3 billion parameters for the generation of node embeddings while using edge features by combining distance, angle, direction and chain information to form a graph representation to predict the binding affinity [16]. PPI-Graphomer uses Evolutionary Scale Modelling 2 (ESM2) and ESM-IF1 to leverage sequence and structural information, respectively, and uses graph neural networks constructed from protein 3D structure for the binding affinity prediction. SSIF-Affinity applies geometric constraints to identify protein binding regions and uses cross-modal attention to combine binding-region structural features and full-length sequence for the binding affinity prediction [21].

Meanwhile, sequence-based models only use protein amino acid residue sequences as input to predict the binding affinity. Such models include ISLAND and ProtT-Affinity [24, 25]. To predict binding affinity, ISLAND uses amino acid composition features, average BLOSUM-62 features, propy features, position-specific scoring matrix features and ProtParam features with kernel representations for its regression model [24]. Meanwhile, ProtT-Affinity uses ProtT5 embeddings with a Transformer encoder to represent interaction between sequences in the protein complex [25, 26]. While structure-based models generally perform better, the requirement of having resolved protein-peptide complex structure is stringent and limits their use cases before a definite peptide drug sequence is determined.

In this study, a sequence-based protein-protein binding affinity predictor, CrossAffinity, was built with the aim of improving the prediction power compared to previous models. The model’s performance was evaluated on different benchmark test sets to compare its prediction capabilities with other established models. The importance of the features and architectures of CrossAffinity was also examined through an ablation study. Lastly, a case study through alanine-scanning mutagenesis and another case study to rank nanobodies were used to uncover the learning patterns of CrossAffinity.

## Methodology

### Dataset for Prediction of Protein-Protein Binding Affinity

Dataset for Prediction of Binding Affinity In the training of the binding affinity training model, the training and validation dataset is the same as those of ProAffinity-GNN, totalling 1740 data points, while excluding those whose information can no longer be obtained using the RCSB PDB data bank application programming interface (API). The dataset consisted of protein complex interacting components and the associated binding affinity values. For the training and validation sets, 5-fold cross-validation was used, and the training and validation sets were randomly split. Test sets to compare with other models were also made by Romero-Molina et al., Qian et al., and Kastritis et al., which consisted of protein sequences highly dissimilar to the training and validation set [16, 19, 27]. The test sets are named S34, S90, S90 subset, S79* and SKEMPI.

### Representation of Protein Complex

The large protein language model, ESM2, was used to generate the features required to build the binding affinity prediction model. The 650 million parameters of the ESM2 model, which generates 1280 dimensions for each residue embedding, were used in this work [28]. The ESM2 model’s attention scores were used by a previous work to predict the probability contact map of a given protein sequence using a logistic regression head [29]. The predicted probability contact map was used in this work to create a graph representation of each protein sequence. Each node represents the amino acid of the protein, and an edge is added between two nodes if the contact probability of the amino acids is more than or equal to 0.5. Each protein sequence was input into the ESM2 model to generate the graph node embedding of each residue. Each protein complex was split into two parts, comprising the binding affinity calculation interface, following the work by Zhou et al [17].

### CrossAffinity Prediction Model Architecture

The dimension of all graph node embeddings was reduced using a multi-layer perceptron (MLP) using equation 1, whereby ***W*** is the weight parameters, *x* is the input, and *σ* is the activation function, before undergoing the graph convolution layer (GCN) to learn the predicted spatial information embedded in the predicted protein contact map. Equation 2 denotes the GCN layer, with ***H***^**(*l*)**^ as the node representations at layer l, ***W*** ^(*l*)^ as the weight parameters at layer *l*, and 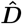 as the diagonal degree matrix. ***Â*** is calculated using the adjacency matrix ***A*** and the identity matrix I as indicated in equation 3.

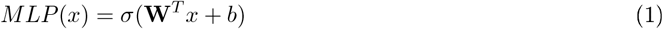

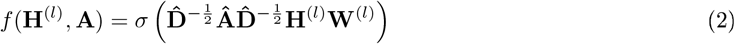

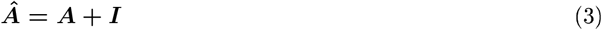

For the protein sequences in both parts to have contextual information of each other, a cross-attention module was implemented as in equation 4. ***Q, K*** and ***V*** are the query, key and value projections of the input token embeddings. ***K*** and ***V*** are from the opposing part of ***Q***. *d*_*k*_ is the number of dimensions of ***K***. The softmax ensures that the attention scores sum up to a total of 1, normalising the contributions of ***V***. A residual connection was also added to include the information from the original token embeddings. Recycling of the cross-attention module was implemented to keep the number of model parameters minimal since the number of data points used to train the model is low.

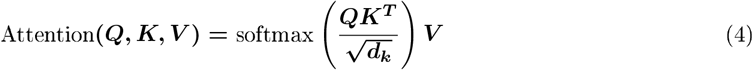

After the cross attention module, the node representations undergo another MLP layer to enrich the information encoded in the node embeddings and graph pooling was performed for each part in the protein complex to obtain summary graph representations. The pooling methods used were max, minimum, mean and attention pooling. Vectors from each pooling method were concatenated. To ensure the invariance of both protein complex parts, the vectors of each protein part were averaged. The last MLP layer aimed to reduce the dimension to 1 to predict the pKd of the protein complex. The pKd of the dataset was calculated as depicted in equation 5.

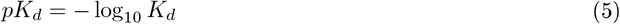

The model was built using PyTorch 2.8.0 and torch-geometric 2.6.1. The analysis was performed using Scipy 1.16.1, Scikit-Learn 1.7.1 and Seaborn 0.13.2.

### Training of the Binding Affinity Model

To ensure a larger penalty and attention for bigger errors, the Huber loss function, as shown in equation 6 with delta equals to 1 and ***n*** equals to the number of data points in a batch, was used to calculate the loss, subsequently averaged, for the backpropagation to train the model parameters. The Adam optimiser was used. A maximum of 150 epochs was set for training. Early stopping criteria were used to prevent overfitting of the trained model by evaluating and comparing the Pearson correlation coefficient performance of the model on the validation set. The training ended if the Pearson correlation coefficient did not improve within 5 epochs.

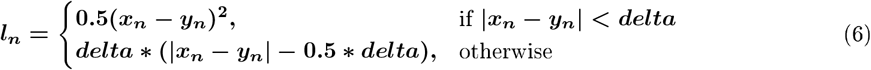

### Model Evaluation Metrics

To evaluate the model’s performance, the Pearson correlation coefficient r, ranging between −1 and 1, was used. It is also the common metric used to compare with other published models. It was calculated from equation 7, whereby ***n*** is the number of data points.

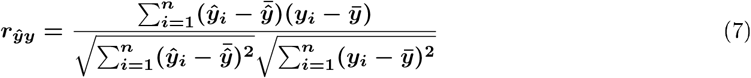

The mean absolute error (MAE) between the actual and predicted binding affinity values is a common metric used in previously published protein-protein binding affinity prediction models and is also used in this work to evaluate and compare the model performance. MAE can be calculated from equation 8.

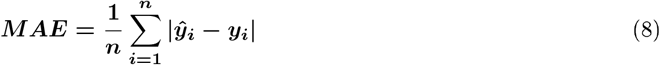

Root mean squared error (RMSE) is also another metric used by some studies to evaluate the model performance. RMSE can be calculated from equation 9.

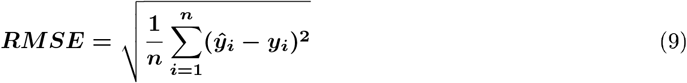

## Results and discussion

### Performance of protein-protein binding affinity on S34 test set

For a fair comparison of performance between CrossAffinity and the other previously published models, benchmark test sets that were prepared by previous studies were used. Test set S34 was prepared by Qian et al., comprising the more recent PDBbind v2021 PPI data, designed to be dissimilar to Qian et al.’s PPAP binding affinity model training set (Table 1) [16]. As the data came from the new entries in the PDBbind v2021 dataset, this ensured that the CrossAffinity model produced in this study, trained using the dataset from the older ProAffinity-GNN model, has yet to come across the data points from this S34 test set. Evaluation of the binding affinity model with **Δ** SASA, which was used as a baseline model in the construction of the PPAP model by measuring the difference in surface area upon protein-protein interaction, revealed that the binding affinity model performs better than the baseline that had an uninformative Pearson correlation coefficient of −0.04. In comparison with the structure-based models, which had richer information of the entire protein-protein binding complex as input, namely ProAffinity-GNN, PPI-Affinity, AAIN_Predictor, PRODIGY and PPAP, the CrossAffinity binding affinity model managed to outperform all these models in all the evaluation metrics. The best performing model, PPAP, had a Pearson correlation coefficient of 0.492, which is lower than the CrossAffinity model trained in this work, which had a Pearson correlation coefficient of 0.575. The MAE and RMSE of CrossAffinity are also shown to be the lowest out of all the other models. This may be due to the large majority of structure-based models being over-trained on their training dataset, memorising the data points and failing to generalise to future unseen data points. Benchmarking against AlphaFold 3 metrics, a generative biomolecule complex structure prediction model, the sequence-based binding affinity model in this work also managed to extract useful information for the prediction of protein-protein binding affinity, which the AlphaFold3 metrics failed to do.

**Table 1:** Performance of protein-protein binding affinity prediction models on S34 test set.

| Model | R↑ | MAE↓<br>(kcal/mol) | RMSE↓<br>(kcal/mol) | Model<br>Type | Remarks |
| --- | --- | --- | --- | --- | --- |
| ProAffinity-GNN [17] | 0.260 | 1.91 | 2.34 | Structure | Model inferenced in this study |
| PRODIGY [18] | 0.138 | 5.33 | 9.88 | Structure | Model inferenced in this study |
| AAIN_Predictor [30] | 0.189 | 2.24 | 2.66 | Structure | Model inferenced in this study |
| PPAP [16] | 0.492 | 1.54 | 1.88 | Structure | Model inferenced in this study |
| ΔSASA [16] | -0.04 | - | - | Structure | Score obtained from PPAP study [16] |
| AF3<sub>ranking_score</sub> [14] | -0.03 | - | - | Sequence | Score obtained from PPAP study [16] |
| AF3<sub>ipTM</sub> [14] | 0.00 | - | - | Sequence | Score obtained from PPAP study [16] |
| AF3<sub>pTM</sub> [14] | -0.22 | - | - | Sequence | Score obtained from PPAP study [16] |
| AF3<sub>pae_interact</sub> [14] | -0.18 | - | - | Sequence | Score obtained from PPAP study [16] |
| **CrossAffinity** | **0.575** | **1.43** | **1.81** | **Sequence** | Model inferenced in this study |

### Performance of protein-protein binding affinity on S90 test set

The S90 test set prepared by Romero-Molina et al. in their PPI-affinity work was also used as an external benchmark dataset to evaluate the binding affinity model created in this work (Table 2) [19]. In this benchmark test set, the CrossAffinity model managed to outperform the structure-based models, PPAP, ProBAN, AAIN_Predictor and PRODIGY, in Pearson correlation coefficient, MAE and RMSE. However, the CrossAffinity model in this work did not outperform ProAffinity-GNN in Pearson correlation coefficient and RMSE. The baseline model, **Δ**SASA, produced in the PPAP model study, which measures the change in surface area upon complex binding, and the AlphaFold3 model also managed to perform better in this test set as compared to test set S34, but was still unable to reach the performance of CrossAffinity. Overall, the structure-based models were able to predict more accurate binding affinity values in this test set as compared to the S34 test set, while the performance of the CrossAffinity model is more stable with fewer changes in the metrics between both test sets. In comparison with ISLAND, another previously published sequence-based model, the CrossAffinity model is better in Pearson correlation coefficient, MAE and RMSE, signifying an improvement in sequence-based models.

**Table 2:** Performance of protein-protein binding affinity prediction models on S90 test set.

| Model | R↑ | MAE↓<br>(kcal/mol) | RMSE↓<br>(kcal/mol) | Model<br>Type | Remarks |
| --- | --- | --- | --- | --- | --- |
| ProAffinity-GNN [17] | 0.652 | 1.53 | 1.98 | Structure | Model inferenced in this study |
| ProBAN [31] | 0.55 | 1.75 | - | Structure | Score obtained from PPAP study [16]<br>Tested on 82 out of 90 data points [16] |
| PRODIGY [18] | 0.313 | 2.54 | 3.61 | Structure | Model inferenced in this study |
| AAIN_Predictor [30] | 0.395 | 1.96 | 2.67 | Structure | Model inferenced in this study |
| PPAP [16] | 0.634 | 1.59 | 2.04 | Structure | Model inferenced in this study |
| ΔSASA | 0.08 | - | - | Structure | Score obtained from PPAP study [16] |
| AF3<sub>ranking_score</sub> [14] | 0.27 | - | - | Sequence | Score obtained from PPAP study [16] |
| AF3<sub>ipTM</sub> [14] | 0.32 | - | - | Sequence | Score obtained from PPAP study [16] |
| AF3<sub>pTM</sub> [14] | 0.49 | - | - | Sequence | Score obtained from PPAP study [16] |
| AF3<sub>pae_interact</sub> [14] | 0.33 | - | - | Sequence | Score obtained from PPAP study [16] |
| **CrossAffinity** | **0.648** | **1.53** | **2.01** | **Sequence** | Model inferenced in this study |

### Performance of protein-protein binding affinity on S90 test subset

As the S90 test set, also known as test set 2 in other studies, consists of data points with IC50, which does not represent protein-protein binding affinity, Zhou et al. removed these data points and formed a subset with 82 entries [17]. In this test subset, all models that were also evaluated on the full S90 test set were noted to have a drop in Pearson correlation coefficient except for AAIN_Predictor, as seen in Table 2 compared to Table 3. Similar to test set S90, the CrossAffinity models managed to outperform some of the structure-based models, PPAP, AAIN_Predictor, PPI-Affinity, PRODIGY, CP_PIE and DFIRE, in both the Pearson correlation coefficient and MAE. However, the CrossAffinity models also did not match the performance of ProAffinity-GNN, PPI-Graphomer and SSIF-Affinity models. Compared to the previous sequence-based models, CrossAffinity showed a massive improvement with a Pearson correlation coefficient of 0.586 compared to that of ISLAND and ProT-Affinity at 0.217 and 0.459, respectively. The MAE and RMSE of CrossAffinity were also lower than those of ISLAND and ProT-Affinity.

**Table 3:** Performance of protein-protein binding affinity prediction models on S90 test subset.

|  |  |  |  |  |  |
| --- | --- | --- | --- | --- | --- |
| Model | R↑ | MAE↓<br>(kcal/mol) | RMSE↓<br>(kcal/mol) | Model<br>Type | Remarks |
| ProAffinity-GNN [17] | 0.635 | 1.44 | 1.88 | Structure | Model inferenced in this study |
| PPAP [16] | 0.558 | 1.56 | 2.01 | Structure | Model inferenced in this study |
| AAIN_Predictor [30] | 0.395 | 1.88 | 2.56 | Structure | Model inferenced in this study |
| PRODIGY [18] | 0.338 | 2.57 | 3.67 | Structure | Model inferenced in this study |
| DFIRE [32] | 0.145 | 26.02 | - | Structure | Score obtained from ProAffinity-GNN study [17] |
| CP_PIE [33] | 0.111 | 7.26 | - | Structure | Score obtained from ProAffinity-GNN study [17] |
| ISLAND [24] | 0.217 | 2.15 | - | Sequence | Score obtained from ProAffinity-GNN study [17] |
| PPI-Affinity [19] | 0.436 | 1.78 | - | Structure | Score obtained from ProAffinity-GNN study [17] |
| PPI-Graphomer [20] | 0.635 | 1.63 | 2.10 | Structure | Score obtained from SSIF-Affinity study [21] |
| SSIF-Affinity [21] | 0.696 | 1.41 | 1.93 | Structure | Score obtained from SSIF-Affinity study [21] |
| ProtT-Affinity [25] | 0.459 | 1.79 | - | Sequence | Score obtained from ProtT-Affinity study [25] |
| **CrossAffinity** | **0.586** | **1.52** | **2.03** | **Sequence** | Model inferenced in this study |

### Performance of protein-protein binding affinity models on S79* test set

Although the S79* test set, synonymous with test set 1 from other studies, which contains two or more protein chains, is derived from a structure-based benchmark database for protein-protein binding affinity and widely used in the evaluation of protein-protein binding affinity models [27]. The test set used in this research omitted 1A2K as it is an obsolete entry in the RCSB PDB databank whose API was used to extract the protein sequence information. Compared to the S90 test subset, models that were evaluated in both test sets, except PPI-Graphomer, have better Pearson correlation coefficient in the S79* test set as seen in Table 3 and Table 4. However, the models in general, except for ProtT-Affinity and AAIN_Predictor, have poorer MAE. In this test set, CrossAffinity performed better than the structural models, AAIN_Predictor, DFIRE, CP_PIE, PPI-Affinity and PPI-Graphomer. However, CrossAffinity underperformed compared to ProAffinity-GNN, PPAP, PRODIGY and SSIF-Affinity.

**Table 4:** Performance of protein-protein binding affinity prediction models on test set S79*.

| Model | R↑ | MAE↓<br>(kcal/mol) | RMSE↓<br>(kcal/mol) | Model<br>Type | Remarks |
| --- | --- | --- | --- | --- | --- |
| ProAffinity-GNN [17] | 0.685 | 1.55 | 2.12 | Structure | *Model inferenced in this study |
| PPAP [16] | 0.658 | 1.68 | 2.18 | Structure | *Model inferenced in this study |
| AAIN_Predictor [30] | 0.581 | 1.87 | 2.46 | Structure | *Model inferenced in this study |
| PRODIGY [18] | 0.73 | - | 1.89 | Structure | †Reported by PRODIGY study [18] |
| DFIRE [32] | 0.602 | 4.64 | - | Structure | †Score obtained from ProAffinity-GNN study [17] |
| CP_PIE [33] | 0.517 | 8.80 | - | Structure | †Score obtained from ProAffinity-GNN study [17] |
| ISLAND [24] | 0.378 | 2.10 | - | Sequence | †Score obtained from ProAffinity-GNN study [17] |
| PPI-Affinity [19] | 0.616 | 1.82 | - | Structure | †Score obtained from ProAffinity-GNN study |
| PPI-Graphomer [20] | 0.622 | 1.69 | 2.24 | Structure | †Score obtained from SSIF-Affinity study [21] |
| SSIF-Affinity [21] | 0.710 | 1.49 | 2.00 | Structure | †Score obtained from SSIF-Affinity study [21] |
| ProtT-Affinity [25] | 0.628 | 1.65 | - | Sequence | †Score obtained from ProtT-Affinity study [25] |
| **CrossAffinity** | **0.644** | **1.67** | **2.17** | Sequence | *Model inferenced in this study |

\* Obsolete PDB ID 1A2K is omitted.
† Evaluated on full S79 or test set 1.

### Performance of protein-protein binding affinity on SKEMPI test set

In many of the previously published protein-protein binding affinity models, the SKEMPI dataset, which contains information on binding affinity changes upon mutations in protein-protein interactions, was used to evaluate the models’ performances. The binding affinity prediction structure-based models, PPI-Affinity, ProAffinity-GNN and PPAP, although not specifically trained to identify changes in binding affinity upon protein mutation, managed to achieve comparable performance to the mutation-specific models, signifying that they managed to learn useful underlying trends for very small changes in amino acid sequences in a protein complex. Likewise, even without the more comprehensive structural information, CrossAffinity managed to achieve comparable performance to the structure-based models. CrossAffinity managed to outperform PPI-Affinity, ProAffinity-GNN and SSIF-Affinity. Although the Pearson correlation coefficient of CrossAffinity is lower than that of PPAP, its performance still came very close to that of PPAP.

### Inference time of the protein-protein binding affinity prediction models

Although the inference time of a model is of little consideration in practical usage with few data points, the accumulated impact when evaluating many data points can be massive. An example of inferencing models on a huge dataset is when using such models to predict a huge library of mutated sequences to identify strong peptide binders to a target protein in a peptide drug discovery effort. Using S34 and S90 test sets to evaluate the models’ speed, CrossAffinity was noted to be exceptionally fast as compared to the other structure-based models, even when using CPU only. CrossAffinity was only slower than PRODIGY on the smaller S34 test set but faster than PRODIGY on the larger S90 test set. The inference speed of CrossAffinity was effectively halved again when the A6000 GPU was used instead.

### Ablation study

The model design of CrossAffinity was examined through ablation of model architectures and features. Upon ablation of either the cross-attention module, the GCN layer, or the ESM2 embeddings, the model performance decreased. This signifies that these components contain important information which the CrossAffinity model uses in order to predict the protein-protein binding affinity. It was noted that the worst performance came when the ESM2 embeddings were swapped with one-hot encoding of the 21 amino acid residues. One possible reason is that the enormous evolutionary information encoded in the pre-trained ESM2 model embeddings, which are learned by training on the ESM2 model on the enormous UniRef50 dataset, is important for protein-protein binding affinity [39]. This transfer learning from the ESM2 model is clearly the most informative feature of CrossAffinity to predict the protein-protein binding affinity. The ablation of cross attention led to a small decrease in performance for S72* but a more observable decrease in S34, S90 and S90 subset test sets. The decrease in performance may be due to the importance of allowing the two parts of the protein complex to exchange and gather residue context of the opposing part so that the information of each residue embedding in the model is enriched. Ablation of the GCN layer also led to worse performance in all the test sets. However, the performance was still better than removing the cross-attention module. This may be due to the fact that the local neighbourhood information within the protein complex part is less important than the contextual information of the interacting protein complex part.

### Case studies

Alanine scanning mutagenesis first started out as a wet-lab technique to identify key residues in the human growth hormone (hGH) that play an important role in its binding to the hGH receptor [40]. The procedure was executed through the replacement of residues by alanine one at a time and observing the impact of each residue mutation on the protein-protein binding affinity [40]. Likewise, this method can also be used *in silica* [41]. Applying alanine-scanning mutagenesis to the yeast Cdc13 and Stn1 complex, certain patterns in the predicted binding affinity were revealed. In Cdc13, the mutation of interface residues was noted to result in lower predicted pK_d_ as compared to the predicted pK_d_ of the original Cdc13 sequence, as seen in Figure 3A. The pK_d_ of these interface residues was also observed to be low compared to the neighbouring residues in sequence, showing that the model could learn the importance of these interface residues. In a global context, these interface residues generally lie in the second half of the Cdc13 sequence and the residues. When the alanine mutation occurs in the second half of the sequence, the predicted pK_d_ was observed to be lower than when the mutation occurs in the first half. When the alanine-scanning mutagenesis was performed on the Stn1 sequence, the predicted pK_d_ of the interface residues were also noted to be lower than the original Stn1 sequence as observed in Figure 3B. Some, but not all, of the interface residues were noted, when mutated to alanine, to result in lower predicted pK_d_ when compared to the neighbouring residues. This shows that although the model might possibly have implicitly learned to identify some of the interface residue, even though it is not perfect, through only the protein sequences, while such information is given explicitly as inputs to the structure-based models.

**Figure 1.**
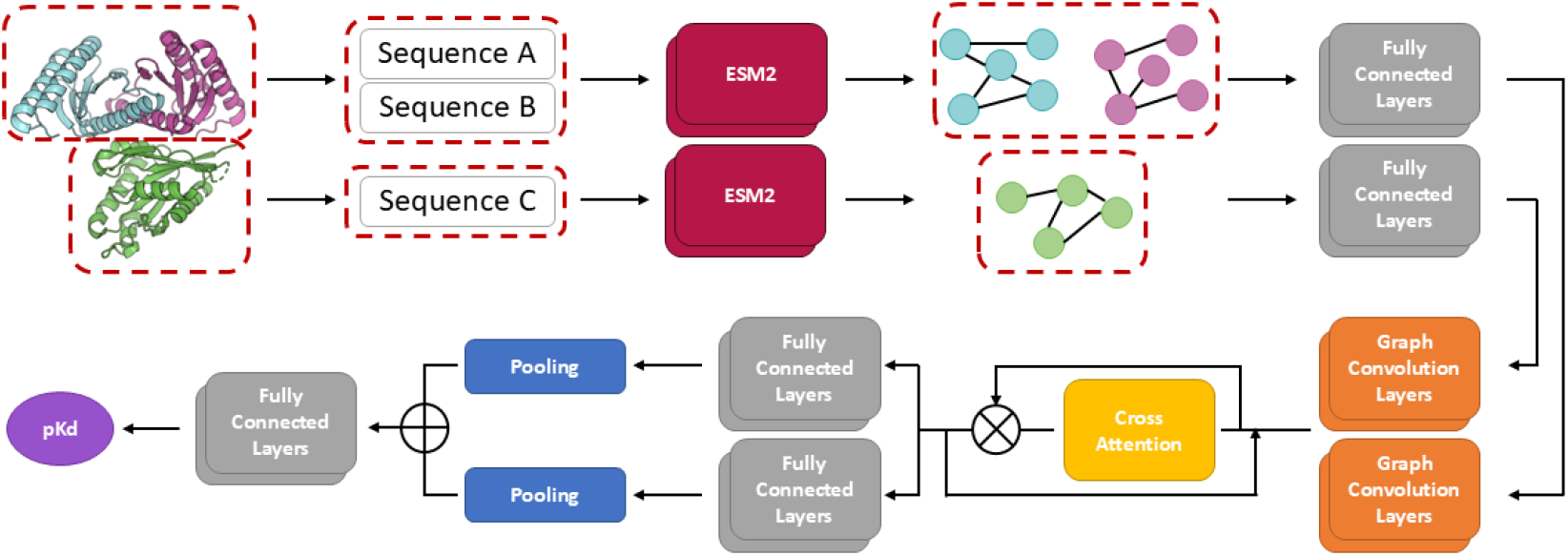
Architecture of sequence-based CrossAffinity protein-protein binding affinity model.

**Figure 2.**
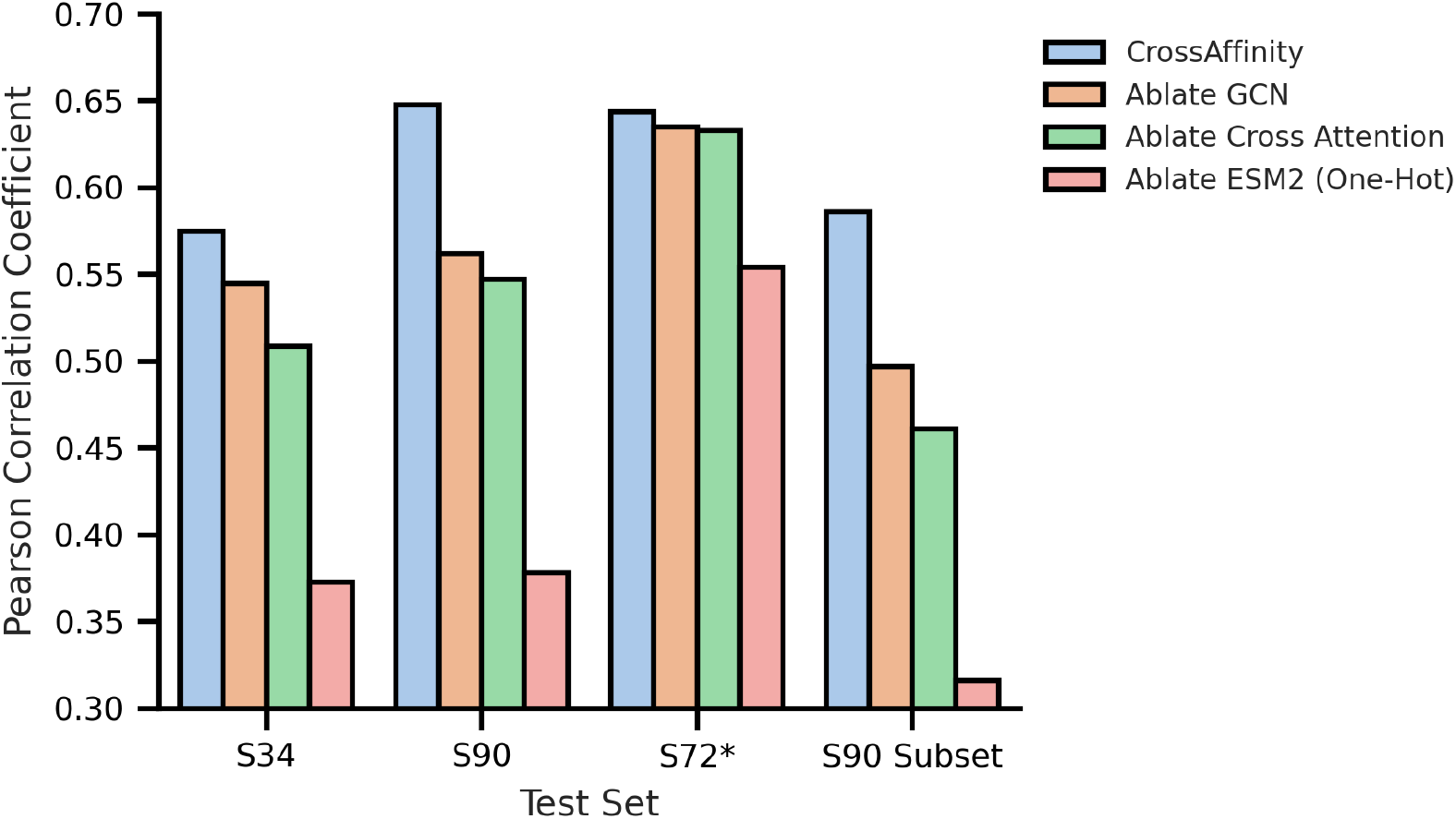
Impact of model features and architectures ablation on CrossAffinity’s Pearson correlation coefficient.

**Figure 3.**
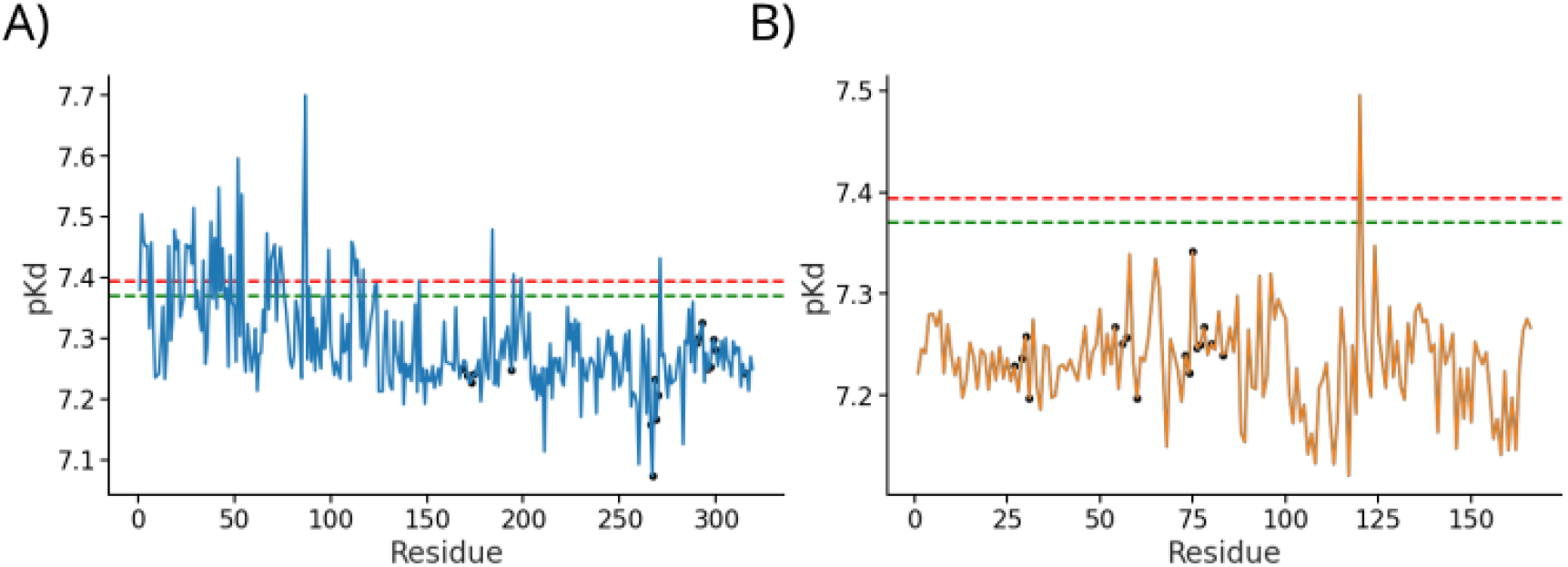
Alanine scanning site-directed mutagenesis on yeast Cdc13 and Stn1. Alanine scanning performed on (A) Cdc13 and (B) Stn1. The red dotted line is the true pK_d_ while the green dotted line is the predicted pK_d_ of the original sequence. The black-filled circles are the residues within 5 Aof the interacting protein.

*Escherichia coli* is a model organism, frequently used for the expression of protein of interest [42, 43]. This is due to the low cost of fermentation, availability of efficient transformation tools and deep understanding of the *Escherichia coli*’s genetics [44]. One method to purify the expressed recombinant protein is immobilised metal affinity chromatography (IMAC), in which a small peptide tag at the N- or C-terminus of the recombinant protein binds to immobilised bivalent metal ions [45]. Upon washing off the non-adhering molecules, the adsorbed recombinant protein can be eluted by lowering the pH [46, 47]. However, there are also endogenous bacterial proteins, such as FKBP-type peptidyl-prolyl cis-trans isomerase (SlyD), which may have an affinity for divalent metal ions, causing them to be co-purified with the recombinant protein of interest as a contaminant [44, 48]. To resolve this issue, Hu et al. identified nanobody sequences with high affinity to SlyD through immunising a dromedary and subsequent phage display and panning [44]. These nanobodies can be used to remove the SlyD contaminant via immunoprecipitation [44]. Through the known binding affinities of these nanobodies obtained using enzyme-linked immunosorbent assay (ELISA) by Hu et al., CrossAffinity performance may be evaluated in ranking these nanbodies’s affinity to SlyD. By analyzing the predicted pK_d_ of the nanobody sequences with SlyD, CrossAffinity was noted to have good performance of 0.776 in Pearson correlation coefficient and 0.800 in Spearman correlation coefficient from the results in Table 7. CrossAffinity was able to isolate the nanobody sequence with the highest binding affinity to SlyD. However, it swapped the rank between the second and third rank sequences and between the fourth and fifth rank sequences. Even so, CrossAffinity could still identify that the second and third rank sequences have better binding affinity than the fourth and fifth rank sequences to SlyD. This shows that CrossAffinity can perform ranking of these nanobody sequences to obtain high-affinity nanbodies for real use scenarios.

**Table 5:**
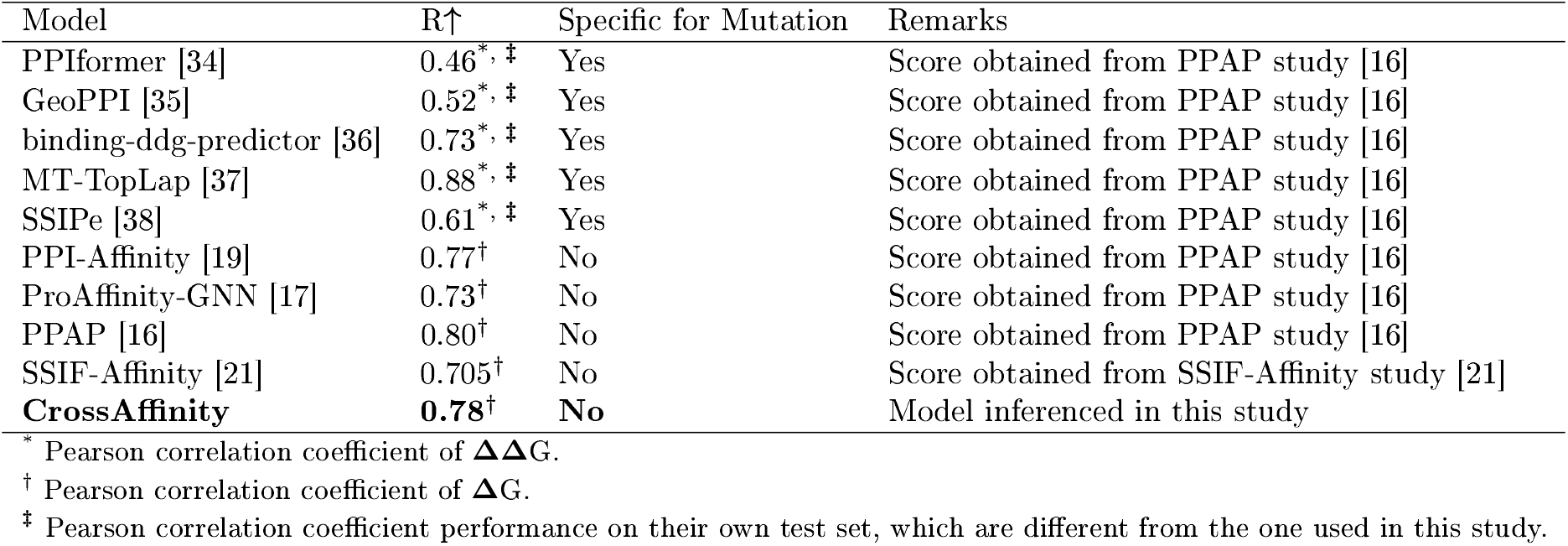
Performance of protein-protein binding affinity prediction models on SKEMPI test set.

**Table 6:** Inference time comparison for different models.

| Model | S34 / seconds | S90 / seconds | Model Type | Remarks |
| --- | --- | --- | --- | --- |
| ProAffinity-GNN | 1106.93 | 2265.6 | Structure | Model inferred in this study |
| PPAP | 896.90 | 1653.7 | Structure | Model inferred in this study |
| PRODIGY | 37.433 | 271.52 | Structure | Model inferred in this study |
| AAIN_Predictor | 234.31 | 350.10 | Structure | Model inferred in this study |
| **CrossAffinity** | **56.097** | **79.967** | **Sequence** | Model inferred in this study |
| **CrossAffinity\*** | **20.793** | **35.192** | **Sequence** | Model inferred in this study |

<sup>\*</sup> ESM2 on A6000 GPU.

**Table 7:** Comparison of experimental and predicted antibody affinity against SlyD.

| Antibody Name | Affinity (nM) | Experimental Ranking | Predicted $pK_d$ | Predicted Ranking |
| --- | --- | --- | --- | --- |
| Nb5 | 1.5 | 1 | 8.5607 | 1 |
| Nb12 | 2.5 | 2 | 8.4917 | 3 |
| Nb21 | 4.5 | 3 | 8.4944 | 2 |
| Nb127 | 25 | 4 | 8.4757 | 5 |
| Nb3 | 32 | 5 | 8.4758 | 4 |

## Conclusion

In this study, CrossAffinity, a sequence-based protein-protein binding affinity prediction model, was constructed. The model uses a graph representation of the predicted contact map of the ESM2 model with node embedding obtained from the ESM2 model as well. The cross-attention module allows the model to understand the contextual information of the interacting components in the protein complex. The model managed to outperform all other structure-based and sequence-based models in the S34 test set, which contains newer protein complex structures and binding affinity values. In other test sets, although CrossAffinity does not perform the best in every single test set, it still managed to outperform all other sequence-based models and some structure-based models. Despite the lesser information as input into CrossAffinity compared to the structure-based models, its performance is still comparable to these structure-based models. Given that the majority of protein complex structures are still undetermined, CrossAffinity, as a sequence-based model, can provide a reasonable prediction in these situations. In the event of isolating top-performing peptide drug sequences from a huge sequence space, the quick inference and the lack of requiring protein complex structure when using CrossAffinity may also speed up the drug discovery efforts.

## Acknowledgements

The authors wish to thank the Singapore Ministry of Education (MOE) for funding and making this study possible through a Tier 1 grant RG97/22.

## Data availability

The codes to train and use CrossAffinity are available at https://github.com/CodeJS19/CrossAffinity.

## Supporting information

- training.xlsx: PDB IDs of the training dataset of CrossAffinity.
- Supplementary_Info.pdf: Figure S1 is the hyperparameter optimisation result of CrossAffinity; Figure S2 is the Pearson correlation coefficient of the best-performing model during the training process.

