## Supplementary Information for "CrossAffinity: A Sequence-Based Protein-Protein Binding Affinity Prediction Tool Using Cross-Attention Mechanism"

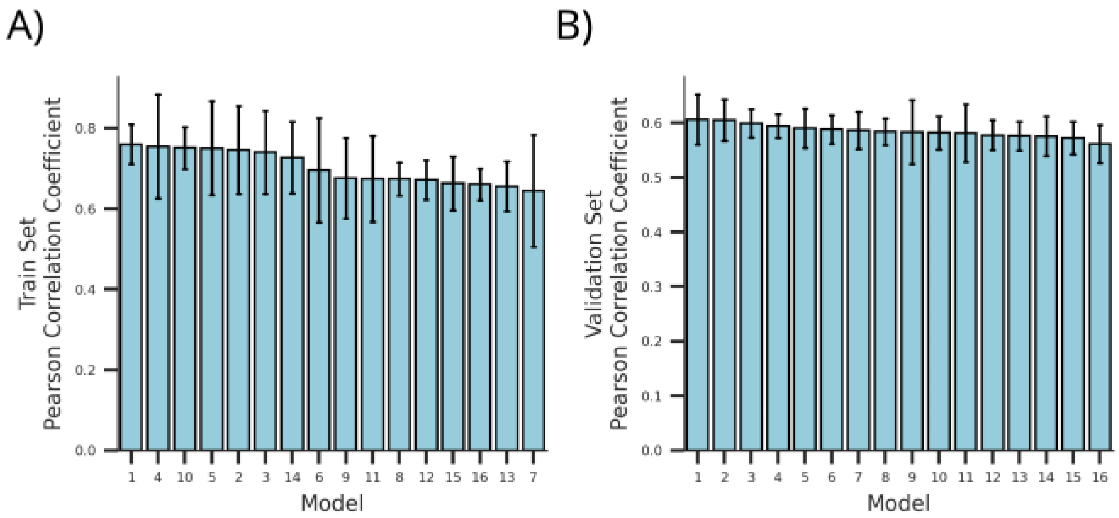


Figure S1: Hyperparameter optimisation of CrossAffinity using Pearson correlation coefficient. Pearson correlation of models on the randomly split (A) training and (B) validation set. Error bars are the standard deviation of the 5-fold cross-validation models. Models are sorted based on best performance on the validation set.

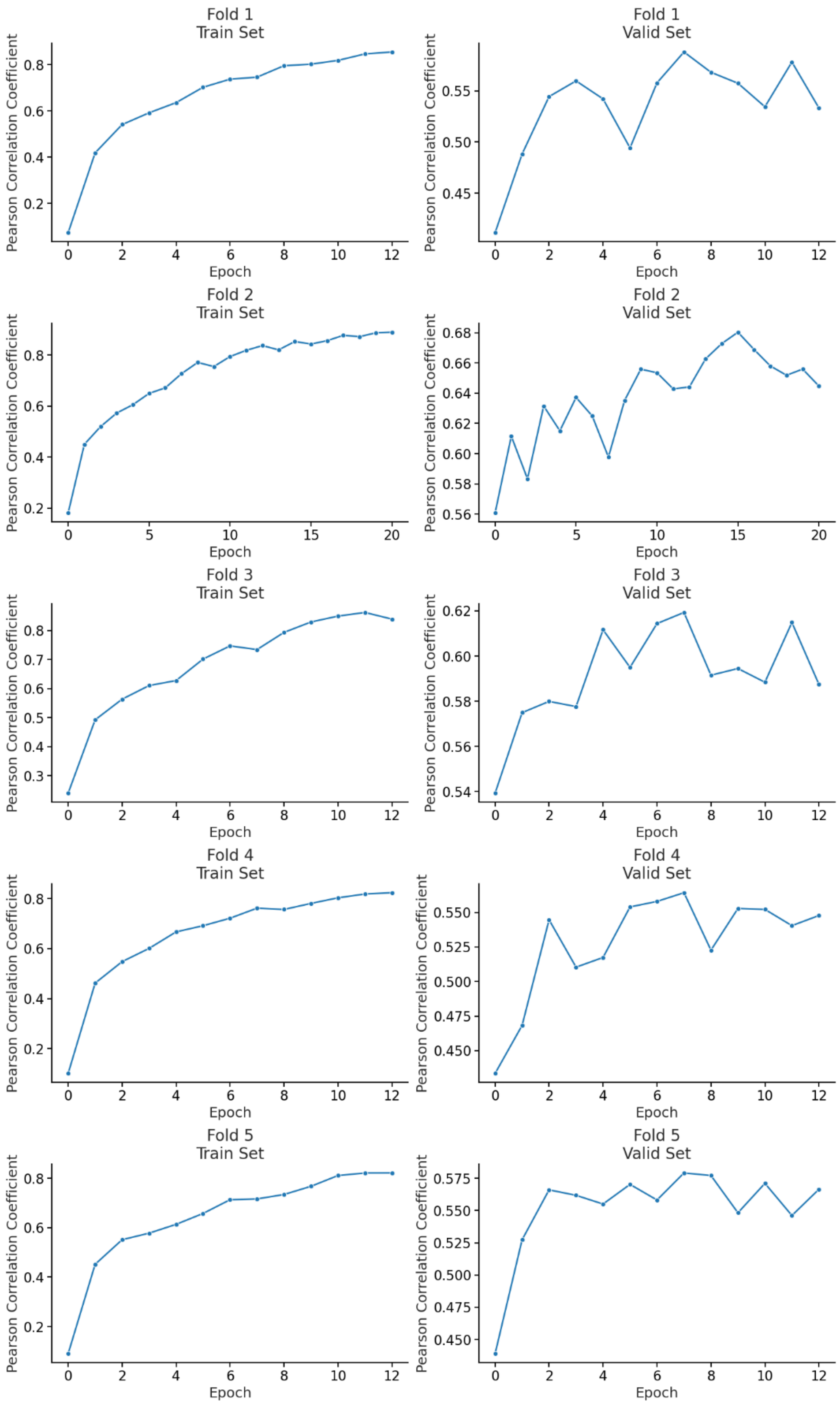


Figure S2: Pearson correlation coefficient of the 5-fold cross-validation models belonging to the best-performing model 1.
